# Single-cell characterization of bacterial optogenetic Cre recombinases

**DOI:** 10.1101/2025.06.06.658346

**Authors:** Hellen Huang, Fereshteh Jafarbeglou, Mary J. Dunlop

## Abstract

Microbial optogenetic tools can regulate gene expression with high spatial and temporal precision, offering excellent potential for single-cell resolution studies. However, bacterial optogenetic systems have primarily been deployed for population-level experiments. It is not always clear how these tools perform in single cells, where stochastic effects can be substantial. In this study, we focus on optogenetic Cre recombinase and systematically compare the performance of three variants (OptoCre-REDMAP, OptoCre-Vvd, and PA-Cre) for their population-level and single-cell activity. We quantify recombination efficiency, expression variability, and activation dynamics using reporters which produce changes in fluorescence or antibiotic resistance following light-induced Cre activity. Our results indicate that optogenetic recombinase performance can be reporter-dependent. Further, single-cell analysis revealed highly heterogeneous activity across cells. Although general trends match expectations for light-dependent recombination, we found substantial variation in the efficiency and timing of recombinase activity from cell to cell. These findings suggest critical criteria for selecting optogenetic recombinase systems and indicate areas for optimization to improve single-cell capabilities of bacterial optogenetic tools.

## Introduction

Optogenetic methods have the potential to enable tunable and precise regulation of gene expression for a myriad of synthetic and systems biology applications. For example, in bacteria, light has been used to control the timing of biochemical synthesis^1^, modulate biofilm formation^2^, and regulate cellular growth^3,4^. However, the majority of bacterial optogenetic studies to-date have focused on population-level regulation^5^. The ability to spatially pattern light offers the enticing potential for single-cell resolution control. Tools that enable the regulation of gene expression in single cells could be used to probe the impact of spatial structure within microbial communities^6–11^, to mimic stochastic events^12–14^, or to drive gene expression in individual bacteria^8,15–17^. For example, light-dependent recombination has been used to model the stochastic deletion of a drug resistance gene^18^ and single-cell control has been used to modulate gene expression dynamics^15,16,19–22^. For these applications, light offers distinct advantages over classical approaches using chemical induction. Light is amenable to single-cell resolution spatial targeting, whereas chemical inducers are challenging to apply spatially and are subject to diffusion^23^. In addition, light is straightforward to add and remove, and because light is orthogonal to many endogenous cellular processes, optogenetic tools can be used to control cellular function with minimal off-target effects^24^. There is excellent potential for precise regulation of gene expression in single cells using optogenetic regulation via spatiotemporal control with light, however the performance of many optogenetic tools in bacteria has not been characterized at the single-cell level.

In this study we focus on optogenetic recombinases. Recombinases are proteins that recognize a specific 30-50 base pair sequence of DNA and mediate site-specific recombination based on the orientation of the recognition sites. For example, Cre recombinase is a widely used tyrosine recombinase derived from P1 bacteriophage that recognizes a 34 base pair site called loxP. Cre will either excise, invert, or translocate the DNA based on the orientation of the loxP sites^25^. Cre recombinase has been used in synthetic biology applications to construct complex cellular logic circuits^26,27^ and to engineer gene circuits with synthetic memory^28^. Light-inducible recombinases can allow specific cells within a population to be targeted or can be used to control the timing of recombination. For cases like DNA excision where the action of Cre is irreversible, once recombination occurs the cell will remain in that state even after light illumination stops, which can be useful for avoiding phototoxicity due to extended light exposure. Several studies have developed light-responsive Cre recombinase variants. A common approach involves splitting the Cre protein and linking it to light-sensitive photodimers. For these systems, light illumination brings the photodimers together, allowing Cre to form a functional enzyme. Optogenetic recombinases have been developed for mammalian cells^26,27,29,30^ and *Saccharomyces cerevisiae*^31,32^. In bacteria, optogenetic Cre variants include the OptoCre-REDMAP^29^ and OptoCre-Vvd^34^ systems that we developed in prior work, and the PA-Cre system introduced by Koganazawa et al.^18^. These systems have been used for a wide range of applications, from regulating antibiotic resistance gene expression^18,33,35^, to improving biofuel production^35^, to capturing spatial information via light as a form of memory^36^. Although these studies have demonstrated the practical utility of optogenetic Cre, they have not focused on single-cell regulation.

We conducted a comprehensive comparison of these three optogenetic Cre systems, quantifying their activities at both the population and single-cell levels. We first evaluated the efficiency of these systems using three distinct reporter types. Our data show that all three systems are functional, but interchanging optogenetic Cre systems and reporters can require additional optimization for robust performance. We next used microscopy and microfluidics to monitor single-cell expression dynamics. Single-cell analysis revealed heterogenous activity, including examples of broad expression distributions or bimodal expression patterns. We also observed significant cell-to-cell variability in expression timing and maximum expression levels. These findings provide a quantitative comparison of bacterial optogenetic Cre recombinase variants and identify constraints and opportunities for optimizing recombinase function in single cells.

## Results

We began by comparing the performance of OptoCre-REDMAP, OptoCre-Vvd, and PA-Cre at the population level. The three systems were developed and characterized using distinct reporters, so our initial goal was to compare the performance of the Cre variants under identical contexts. OptoCre-REDMAP is red light responsive^33^ and consists of two split Cre fragments attached to FHY1 and ΔPhyA photoreceptors derived from *Arabidopsis thaliana*^30,37^ (Fig. 1A). ΔPhyA binds to FHY1 under red light (∼660 nm) and dissociates under far-red light (∼730 nm). The two split Cre fragments are located on an operon controlled by the IPTG-inducible P_lacUV5_ promoter. The REDMAP design requires the chromophore phycocyanobilin (PCB) and we included the gene encoding it (*ho1-pcyA*) on the plasmid with OptoCre-REDMAP (Fig. S1). OptoCre-Vvd has a similar design^34^, consisting of split Cre fragments fused to Vivid (Vvd) photodimers derived from the fungus *Neurospora crassa*^38^. The Vvd fragments homodimerize under blue light (∼465 nm) and dissociate in the dark^39^. As with OptoCre-REDMAP, the optogenetic construct is encoded on an operon under the control of P_lacUV5_. In contrast to REDMAP, the chromophores required for Vvd are flavin nucleotides, which are endogenous to all bacteria^24^, and thus it was not necessary to add genes encoding them. PA-Cre is a blue light responsive optogenetic Cre system^40^ that was adapted for bacteria^18^. It consists of split Cre fragments attached to the magnet photodimers pMag and nMag, which are engineered variants of Vvd^41^. Blue light (∼465 nm) causes pMag and nMag to heterodimerize and they dissociate in the dark. The genes encoding PA-Cre are present on an operon that is controlled by the IPTG-inducible P_LlacO1_ promoter, which is weaker than P_lacUV5_^42^. All three optogenetic Cre systems use the same split site, where the nCre fragment consists of the first 43 amino acids of Cre, with cCre containing the remainder.

**Figure 1.**
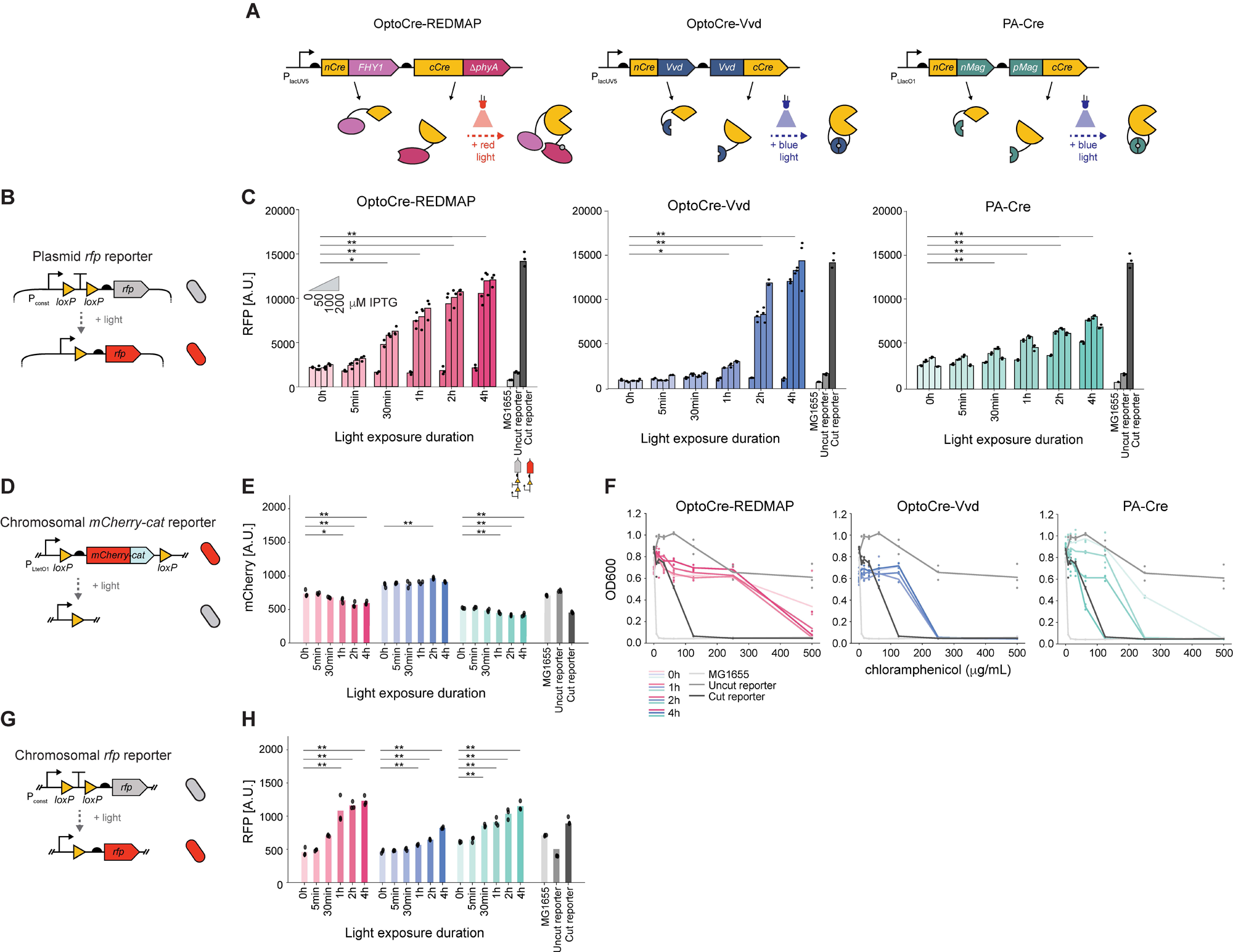
Population-level comparison of three optogenetic Cre systems on three reporters. **(A)** Schematics of the three optogenetic Cre systems: OptoCre-REDMAP, OptoCre-Vvd, and PA-Cre. In all cases, the recombinase is split into nCre and cCre fragments, which are fused to photodimers. Small gray circles indicate chromophores and correspond to phycocyanobilin (PCB) for OptoCre-REDMAP and flavin for OptoCre-Vvd and PA-Cre. The ribosome binding site is represented by a black semicircle upstream of the coding sequence. **(B)** Plasmid *rfp* reporter contains a constitutive promoter followed by a loxP-flanked terminator in front of the *rfp* gene. As shown in the schematic, cells are expected to be dark in the uncut state and to express RFP in the cut state. The terminator is represented by a “T” symbol. **(C)** Population-level fluorescence data comparing the performance of the three optogenetic Cre variants on the plasmid *rfp* reporter. For each light exposure duration, we tested four IPTG concentration 0, 50, 100, and 200 µM. Controls include wild type *E. coli* MG1655 with no additional constructs, an uncut version of the reporter, and a cut version of the reporter. A.U., arbitrary units. Bars represent the mean values, with individual biological replicates shown as dots (n = 3). Statistical significance was determined by one-way ANOVA followed by Tukey’s post-hoc test. *, *p* < 0.05; **, *p* < 0.001. One-way ANOVA for the 100 µM case is shown; for other IPTG concentrations see Fig. S2. **(D)** Chromosomal *mCherry-cat* reporter contains a translational fusion of *mCherry* and *cat* flanked by loxP sites. With no recombinase activity, cells are expected to express mCherry and be resistant to chloramphenicol; after excision cells are expected to have low fluorescence and exhibit chloramphenicol sensitivity. **(E)** Population-level experiments comparing fluorescence for strains with the three optogenetic Cre variants and the chromosomal *mCherry-cat* reporter. Experiments use 100 µM IPTG induction. Bars represent the mean values, with individual biological replicates shown as dots (n = 3). Statistical significance was determined by one-way ANOVA followed by Tukey’s post-hoc test. *, *p* < 0.05; **, *p* < 0.001. **(F)** Antibiotic resistance data comparing the optical density at 600 nm (OD600) at different chloramphenicol concentrations and light exposure durations. Individual replicate data are plotted as dots. **(G)** Chromosomal *rfp* reporter contains a loxP-flanked transcriptional terminator in front of *rfp*. **(H)** Population-level measurements comparing the RFP level for the three optogenetic Cre systems acting on the chromosomal *rfp* reporter. Experiments use 100 µM IPTG induction. Bars represent the mean values, with individual biological replicates shown as dots (n = 3). Statistical significance was determined by one-way ANOVA followed by Tukey’s post-hoc test. *, *p* < 0.05; **, *p* < 0.001.

To directly compare the functionality of the optogenetic Cre systems, we tested all three designs using the same reporters. First, we used a plasmid-based reporter that contains a constitutive promoter followed by a loxP-flanked transcriptional terminator upstream of the gene encoding red fluorescent protein (*mrfp1*, denoted hereafter as *rfp*) (Fig. 1B). In the dark, the terminator prevents expression of RFP. When cells are exposed to light, if Cre is functional it will excise the terminator leaving a single loxP site behind and allowing for expression of RFP. We used this plasmid *rfp* reporter when developing both OptoCre-REDMAP and OptoCre-Vvd in previous studies^33,34^, thus we expected these two systems to show robust fluorescence activation in response to light.

Because all three optogenetic Cre systems are controlled by LacI-regulated promoters, expression levels can be modulated by the addition of IPTG. For initial characterization experiments, we tested the three optogenetic Cre systems under a range of IPTG concentrations and light exposure durations, conducting bulk culture experiments in 96 well plates. We induced cells with either 0, 50, 100, or 200 µM IPTG and exposed cells to light for either 0 h, 5 min, 30 min, 1 h, 2 h, or 4 h. For light exposure durations shorter than 4 h, cultures were maintained in the dark until their designated light exposure period, ensuring all cultures finished their light exposure simultaneously. At the conclusion of this period, cultures were refreshed and grown overnight, then RFP levels were quantified using a plate reader.

With the plasmid *rfp* reporter, we found that all three optogenetic Cre systems produced clear increases in RFP expression that were dependent upon both IPTG and light (Fig. 1C, Fig. S2). To establish the expected upper and lower levels of RFP expression, we included several controls. Wild type *E. coli* MG1655 cells establish the lower limit and correspond to autofluorescence levels for our experiments. As an additional negative control, we included cells with just the plasmid reporter and no optogenetic Cre (uncut reporter). In the uncut state, RFP expression should be low due to the presence of the transcriptional terminator. As a positive control, we constructed a reporter plasmid containing only one loxP site and no terminator, mimicking the result of successful Cre recombination (cut reporter). Both OptoCre-REDMAP and OptoCre-Vvd were able to reach RFP expression levels similar to the cut reporter after 4 h of light exposure, while PA-Cre reached about half this level. PA-Cre also exhibited leaky cutting, even in the absence of IPTG, displaying substantial activation of RFP after 4 h of light exposure for the 0 µM IPTG condition. Although both OptoCre-REDMAP and OptoCre-Vvd reliably induced RFP expression with 1-2 h of light exposure, OptoCre-REDMAP produced clear differences in RFP with shorter exposure durations, and we saw increases in RFP expression in response to 5-30 min of light with this system. In general, we found that induction of Cre with IPTG was necessary to achieve full expression of RFP, with 50 µM IPTG or higher leading to substantial expression of RFP.

Since the OptoCre-REDMAP and OptoCre-Vvd constructs are controlled by a different promoter than the PA-Cre design (P_lacUV5_ vs. P_LlacO1_) and the ribosome binding site differs between the designs, we asked whether these distinctions were responsible for the differences in function we observed. To test this, we created a variant of PA-Cre, using the same promoter and ribosome binding site as the OptoCre-REDMAP and OptoCre-Vvd designs (Fig. S3A). These changes resulted in maximal levels of RFP expression that were comparable to the cut reporter positive control, with less leakiness in the 0 µM IPTG state. However, the system exhibited only low dependence on light exposure, with substantial RFP expression even in the dark condition (Fig. S3B). It is possible that further tuning of IPTG concentrations could identify conditions with better light-responsiveness, but these results underscore that the three optogenetic recombinases have distinct patterns of light-induced activation.

Although the original PA-Cre design showed suboptimal activity with the plasmid *rfp* reporter relative to OptoCre-REDMAP and OptoCre-Vvd, this system was developed and optimized for a very different use case that focused on recombination on the chromosome. In the study by Koganezawa et al.^18^ where it was first adapted for bacterial use, PA-Cre was used to excise a chromosomally-integrated reporter with loxP sites flanking mCherry translationally fused with the gene encoding chloramphenicol acetyltransferase (*mCherry*-*cat*) (Fig. 1D). CAT is an enzyme that inactivates chloramphenicol, preventing it from binding to the ribosome, resulting in chloramphenicol resistance^43^. Thus, in the dark, uncut state cells are expected to express mCherry and be chloramphenicol resistant. Light exposure excises *mCherry-cat*, thus the expectation for the cut state is antibiotic sensitive cells with low fluorescence.

We tested the three optogenetic Cre variants with the chromosomal *mCherry-cat* reporter, using identical conditions for the three systems to allow for a direct comparison. We tested a range of light exposure durations and used 100 µM IPTG induction. We chose this IPTG level because all three original studies used this concentration to induce production of Cre^18,33–35^ and our results with the plasmid *rfp* reporter suggest that it is an appropriate concentration for Cre induction (Fig. 1C). We found that mCherry levels showed a modest decrease for the OptoCre-REDMAP and PA-Cre systems (Fig. 1E). Results with OptoCre-Vvd did not show a clear change in mCherry levels.

In addition to measuring mCherry, we also measured cell growth as a function of chloramphenicol to determine light-induced antibiotic resistance levels. This involved refreshing the cells post-light exposure and growing them in a range of chloramphenicol concentrations. The more resistant the cells, the higher the chloramphenicol concentrations they can survive, therefore we expected survival to decrease with increasing light exposure durations. We observed a very slight decrease in survival with increasing light exposure for OptoCre-REDMAP that occurred between the 0 and 1 h light exposure conditions but did not change with additional light. With OptoCre-Vvd, we observed no changes in chloramphenicol resistance, regardless of light exposure duration. PA-Cre showed the clearest response, with a decrease in resistance with increasing light exposure, consistent with *mCherry-cat* excision.

These results demonstrate that the published optogenetic Cre recombinase constructs are optimized for the specific reporters they were designed to work with and that swapping Cre and reporter pairs can lead to suboptimal outcomes without further optimization. Further, these interactions are not always intuitive. For example, the results with the plasmid *rfp* reporter suggest that OptoCre-Vvd cuts more effectively than PA-Cre, however results with the chromosomal *mCherry-cat* reporter show the opposite, with incomplete cutting by OptoCre-Vvd. This is surprising because the Cre variants all target the same loxP sites and, in principle, should be interchangeable, though their cutting efficiencies may vary.

We next asked whether it was possible to develop a reporter that would work with all three optogenetic Cre designs. Additionally, we were interested in ruling out the possibility that elements of the plasmid versus chromosomal reporter designs were responsible for the differences in functions we observed. We integrated our plasmid *rfp* reporter into the *E. coli* chromosome, preserving the constitutive promoter and loxP-flanked terminator design (Fig. 1G). Testing conditions with 100 µM IPTG and a range of light exposure durations, we found that all three optogenetic Cre systems produced an increase in fluorescence with increasing light exposure for the chromosomal *rfp* reporter (Fig. 1H). These results provide additional evidence that all three optogenetic recombinases are functional and they can be used to target the same reporter construct, however the specifics of the design are important for proper function. Further, the relative efficiencies of the recombinases are different and are dependent on the reporter with which they are paired.

To rule out the possibility of phototoxicity due to light exposure and test for any differences in cell viability that were dependent on the recombinase or reporter variants, we also measured growth for each of the optogenetic Cre and reporter pairings (Fig. S4). Light exposure duration had minimal impact on cell growth, suggesting that phototoxicity is not a concern with the light levels we applied for these experiments. We observed weak IPTG-dependent growth impacts with OptoCre-Vvd, but minimal differences for OptoCre-REDMAP or PA-Cre. PA-Cre cultures had slightly reduced growth relative OptoCre-REDMAP and OptoCre-Vvd, but these growth differences were light and IPTG-independent. Thus, recombinase and reporter variants do not introduce a major expression-dependent burden to the cell, though there are modest impacts for some of the variants and IPTG levels.

Population-level measurements provide information about average fluorescence changes in response to light, however it is also useful to understand how single cells respond. For example, applications that capitalize on the spatial targeting of light may be used to induce optogenetic Cre in individual cells. Population-level data can also obscure features such as bimodality or broad variation in fluorescence distributions, which can only be assessed with single-cell resolution measurements. To compare the performance of the three optogenetic Cre systems at the single-cell level, we conducted experiments where we removed samples from the 96 well plates that we had used for the population-level measurements and placed them on agarose pads, using microscopy to quantify fluorescence in single cells.

Cell-to-cell variation can come from several sources, and we were first interested in establishing baselines for our system to distinguish effects like reporter copy number variation from optogenetic Cre-specific effects. Focusing on the controls, we observed that MG1655 cells had uniformly low autofluorescence values (Fig. 2A). The uncut plasmid *rfp* reporter had slightly elevated fluorescence and we observed a distribution of values across cells. Although this slight increase in mean was visible in population-level measurements, single-cell measurements reveal the extent of cell-to-cell heterogeneity. Increases in fluorescence can be attributed to spurious read through of the transcriptional terminator, resulting in a low but detectable level of fluorescence. It is notable that in a small fraction of cells at the upper end of this distribution, expression from the uncut reporter can reach levels comparable to those of the lowest cells in the cut reporter state.

**Figure 2.**
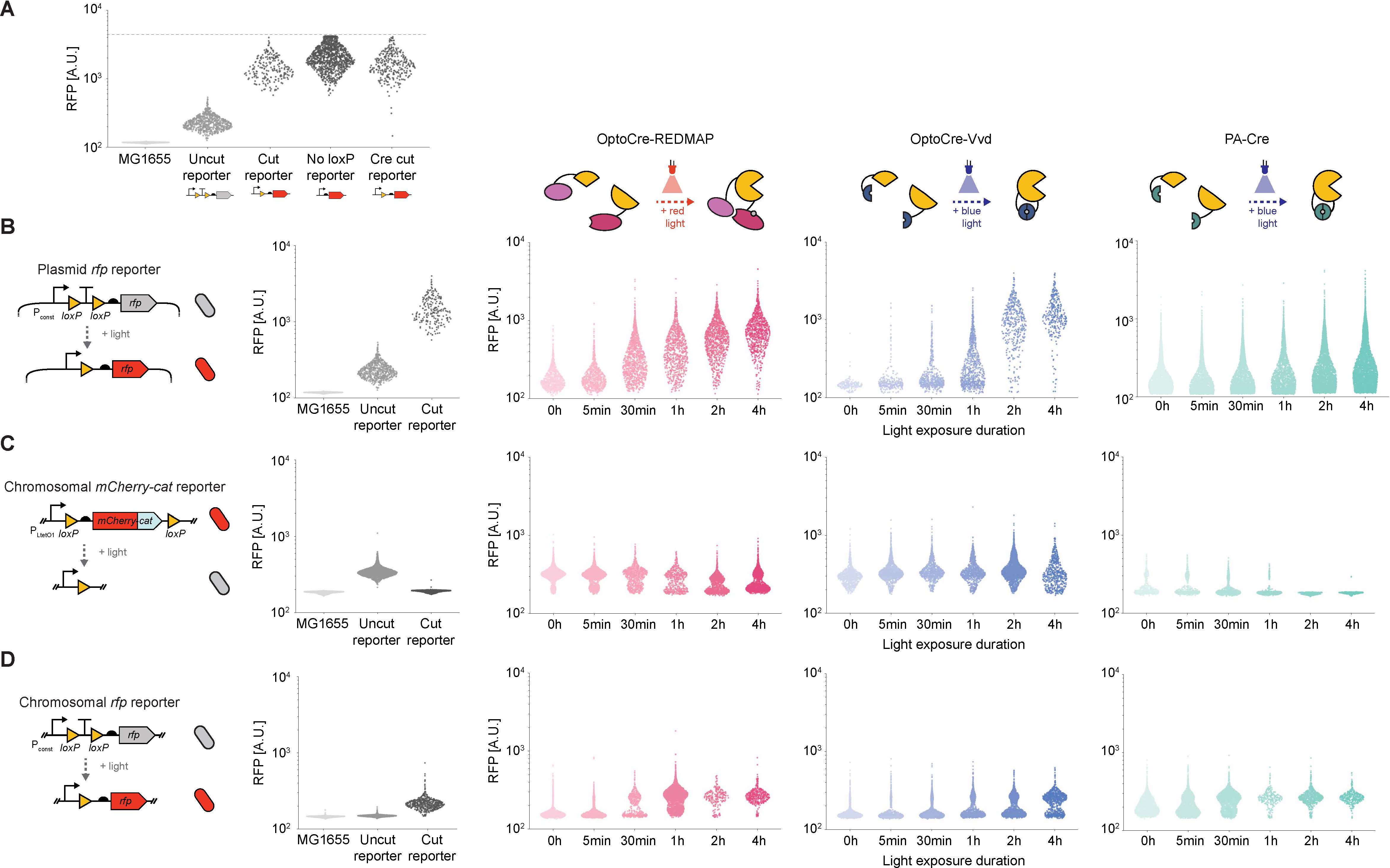
Single-cell resolution comparison of three optogenetic Cre systems on three reporters. **(A)** Control experiments showing violin plots comparing single-cell fluorescence. Data points represent fluorescence data from individual cells. The strains tested are *E. coli* MG1655 with no additional constructs, the uncut plasmid *rfp* reporter, and the cut plasmid *rfp* reporter. In addition, we tested cells with the constitutive promoter driving *rfp* expression with no loxP site, and a cut version of the plasmid *rfp* reporter created by co-transforming the reporter plasmid with a non-optogenetic, non-split version of Cre. Dashed line represents the upper limit of detection using the same imaging settings as the remaining panels in this figure. **(B-D)** Violin plots comparing single-cell fluorescence for strains with the three optogenetic Cre systems acting on the: (B) plasmid *rfp* reporter, (C) chromosomal *mCherry-cat* reporter, and (D) chromosomal *rfp* reporter. All experiments use 100 µM IPTG induction. A.U., arbitrary units.

The cut reporter control showed a clear increase in mean fluorescence over the uncut reporter (Fig. 2A). We were interested in asking whether the presence of a loxP site between the promoter and fluorescent reporter gene introduced heterogeneity in expression, so we constructed an additional positive control where we used the same constitutive promoter to control expression of *rfp*, but omitted the loxP site. Removing the loxP site does increase fluorescence slightly, but distributions are similar between the positive controls with and without the loxP site, suggesting that this is not a major contributor to expression heterogeneity. We also asked whether cutting by Cre could introduce variation in fluorescence. To test this, we co-transformed the plasmid reporter with a non-optogenetic version of Cre that was fully intact (not split). Single-cell fluorescence distributions from these cells were very similar to the cut reporter control. Together, these results suggest that the heterogeneity we observe in the cut reporter is due to factors such as plasmid copy number variation or intrinsic sources of noise, and not differences in transcription introduced by the presence of the loxP site or stochasticity in Cre cutting.

We next quantified the performance of the three optogenetic Cre systems on each of the three reporters in single cells. For these experiments we used six light exposure times: 0 h, 5 min, 30 min, 1 h, 2 h, and 4 h. As with the population-level studies, we used 100 µM IPTG induction. After light exposure, cultures were refreshed and grown overnight, then diluted into fresh media and grown for 2 hours to bring them back to log phase before imaging. Mean results from single-cell microscopy analysis were highly consistent with the population-level findings that we measured using a plate reader. For the plasmid *rfp* reporter, all optogenetic Cre systems produced clear increases in RFP expression (Fig. 2B, Fig. S5). However, single-cell resolution measurements show interesting variation that emerges across cells that was obscured in the population-level measurements. For example, for all systems, we observed very broad distributions in fluorescence, with a few cells exhibiting high fluorescence despite short light exposure durations, and some cells exhibiting low fluorescence despite long exposure periods. This has important implications for the precision of optogenetic induction in single cells because individual bacteria may deviate substantially from mean behavior.

We next performed similar experiments for the Cre variants using the chromosomal *mCherry*-*cat* reporter. The single-cell measurements of OptoCre-REDMAP show evidence of bimodal distributions, likely representing cut and uncut versions of the reporter, with cells shifting to a lower fluorescence state with increased light exposure (Fig. 2C). For OptoCre-Vvd, the single-cell results mirror the population-level data but begin to show trends towards decreasing fluorescence at 4 h of light exposure. For PA-Cre, fluorescence distributions for the 0 h condition are bimodal, however there is a heavy initial bias towards the low state, even without light induction. As light exposure duration increases, cells further shift towards the low state. Although Cre recombinase activity should result in a binary outcome (excised or not excised), comparing results with the plasmid and chromosomal reporters suggests that copy number effects, which are more significant with the plasmid *rfp* reporter construct than the chromosomal reporters, can blur these distributions.

We next looked at single-cell results for the recombinase variants with the chromosomal *rfp* reporter. All three optogenetic Cre systems were able to effectively excise the terminator from the chromosomal reporter, as evidence by the increasing number of single cells with high RFP expression in response to light (Fig. 2D). Again, we observed bimodal distributions, with light-induced Cre shifting cells from a low fluorescence state towards a high fluorescence state at levels commensurate with light exposure. It is interesting to note that the number of cells in the 0 h condition for PA-Cre relative to the other optogenetic Cre systems is higher. These data suggest that PA-Cre exhibits some leaky cutting in the dark state, an effect we also observed with the plasmid *rfp* and chromosomal *mCherry-cat* reporters. To test whether this effect was dependent on IPTG, we conducted single-cell experiments for PA-Cre using the plasmid *rfp* reporter with and without IPTG in the dark state. RFP expression was comparable for the 0 and 100 µM IPTG conditions (Fig. S6), suggesting that this effect cannot be mitigated by adjusting induction levels. Overall, the single-cell resolution results show clear evidence of light-dependent cutting, alongside substantial cell-to-cell heterogeneity.

Although RFP measurements provide valuable insight into variation, they represent the end result of a process that involves multiple steps including DNA excision, then transcription and translation. To better understand where heterogeneity arises, we used a combination of colony PCR and qPCR to assess DNA excision. For these experiments, we focused on OptoCre-REDMAP using 100 µM IPTG. We selected this pairing due to its robust performance across our population and single-cell assays. First, we performed colony PCR experiments to develop a qualitative understanding of DNA excision. After light illumination, cells were diluted and plated on LB agar plates and incubated overnight, allowing single cells to develop into colonies. We performed colony PCR using primers that amplify the region around the loxP sites and used the size of the bands as a qualitative determinant of whether cutting had occurred or not. We performed this experiment on OptoCre-REDMAP with all three reporter types for cultures receiving 0 h or 4 h of light exposure. For each condition, we selected 5 random colonies for colony PCR. Even after 4 h of red light exposure, we found that OptoCre-REDMAP produced incomplete cutting, as indicated by the presence of both the uncut and cut bands in the PCR (Fig. S7). Result with the chromosomal reporters were less variable than the plasmid reporter but we still observed a mixture of outcomes, including complete non-excision, complete excision, or a combination of both uncut and cut state.

In addition to colony PCR, we performed qPCR on cells recovered after 0 h or 4 h of light exposure, focusing on OptoCre-REDMAP with the plasmid *rfp* reporter. After light illumination, cells were refreshed and the plasmid was purified the next day for qPCR analysis. By using primers that bind to the uncut junction, we quantified the amount of uncut plasmid *rfp* reporter DNA remaining after light exposure (Fig. S8). To avoid any spurious issues with DNA quantification that might arise from using controls that contain only one plasmid, we introduced a plasmid containing a catalytically inactive version of Cre (Cre K201A^44^, denoted dCre) alongside our uncut and cut reporters for this analysis. By comparing the amount of uncut plasmid remaining relative to the uncut control, we determined that there was 23.9% DNA recombination in the 0 h condition, compared to 79.9% in the 4 h condition. These results are consistent with our earlier analysis showing that recombination occurs and is light dependent, but recombination is not fully complete even after 4 h of light exposure.

In the previous population-level and single-cell experiments we used a protocol that was conservative about capturing all light-induced gene expression where we completed light exposure, then cultured cells overnight before assessing fluorescent reporter expression. This protocol ensures that Cre recombination, gene expression, and protein maturation have ample time to proceed, allowing us to evaluate the upper limit of expression changes. However, for many single-cell applications the timing of gene expression is important and thus we also conducted studies using an alternative protocol. Here, we assessed fluorescence immediately following light exposure and at subsequent times up to and including the overnight analysis (Fig. 3A). Our goal with this series of experiments was to determine how fluorescence distributions develop across populations of single cells in the period following light exposure.

**Figure 3.**
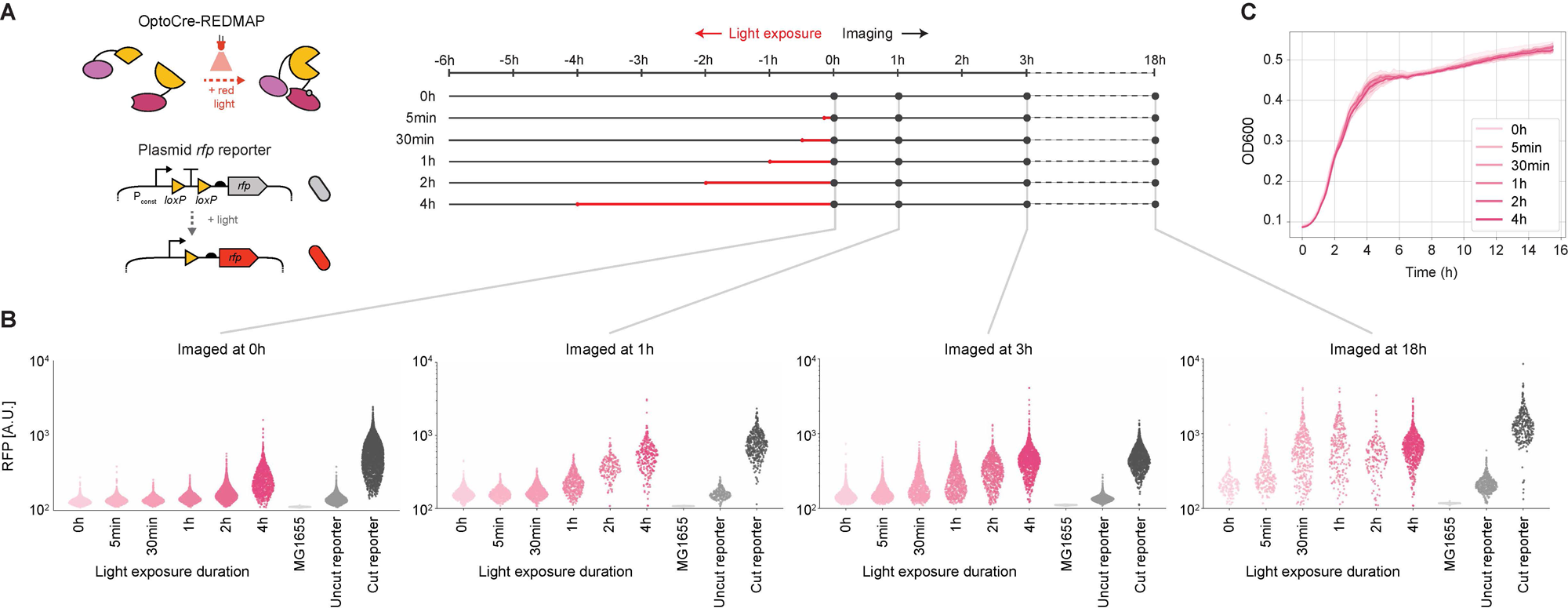
Temporal development of fluorescence distributions in single cells after light exposure. **(A)** Schematic illustrating timepoints for the light exposure and imaging protocols. **(B)** Violin plots comparing single-cell fluorescence measurements conducted at different timepoints following the end of light exposure. All experiments use OptoCre-REDMAP and the plasmid *rfp* reporter with 100 µM IPTG induction. Individual points in the violin plot correspond to single-cell fluorescence data. A.U., arbitrary units. **(C)** Optical density at 600 nm (OD600) over time for cultures where t = 0 h corresponds to the end of light exposure as shown in (A). Error bars show standard deviation around the mean of n = 3 biological replicates.

Again, we focused on OptoCre-REDMAP with the plasmid *rfp* reporter. As before, cultures were induced with 100 µM IPTG for 2 h and then subjected to a range of light exposure durations (0 h, 5 min, 30 min, 1 h, 2 h, 4 h). Cultures were then diluted 1:100 into fresh medium. We then captured single-cell fluorescence using microscopy at 0, 1, 3, and 18 h post-light exposure. We found that fluorescence distributions evolved with time and took many hours to settle to a stable final distribution (Fig. 3B). For example, although 30 min of light exposure was sufficient to produce a fluorescence distribution showing robust activation when imaged at 18 h, expression was modest when imaged 0, 1, and 3 h after light exposure. For reference, the maturation time (t_90_) of RFP is ∼1 hour^45^, therefore a delay of this scale cannot be attributed to maturation of the fluorescent protein alone. Other light exposure durations were similar, requiring many hours to achieve the ultimate fluorescence distributions. We confirmed that cells were growing well following light exposure and that light induction duration had minimal impact on growth (Fig. 3C).

Some of the differences in the fluorescence distributions at 18 h relative to the earlier imaging time points are not due to Cre, as the cut reporter positive control (which does not contain Cre) exhibits higher mean fluorescence at 18 h than at earlier timepoints. Thus, the experimental protocol for this final imaging timepoint, which involves overnight growth (16 h) and a 2 h recovery after dilution in fresh media, leads to higher mean fluorescence. However, this effect does not fully explain the delays we observed in establishing the fluorescence distributions, suggesting that Cre recombination and subsequent gene expression may be heterogeneous following light exposure.

The single-cell snapshot results raise a key question: over what timescale does RFP expression change in an individual cell after light illumination? Although snapshots provide a comprehensive view of distributions of fluorescence across many cells, they do not show how fluorescence changes in an individual cell over time. For example, it is not clear if fluorescence levels slowly increase in all cells together or if individual cells switch to high fluorescence expression, but the timing of this switch varies among cells.

We next aimed to observe the activation dynamics of optogenetic Cre in individual cells. For these experiments, we grew cells containing OptoCre-REDMAP and the plasmid *rfp* reporter in the mother machine microfluidic device^46^ under red light illumination. The mother machine constrains cells to grown in dead-end chambers, enabling continuous imaging of the same cell over many hours of growth (Fig. 4A). After an initial equilibration period where we grew cells in the device in the dark for 2 h, we then illuminated the device with red light for the remainder of the experiment. In addition to uncut and cut reporter controls, we tested OptoCre-REDMAP with 0 or 50 µM of IPTG induction. These two induction conditions allowed us to compare RFP expression in systems with and without OptoCre-REDMAP induced in the same experiment, allowing us to conduct single-cell fluorescence measurements before, during, and after light illumination.

**Figure 4.**
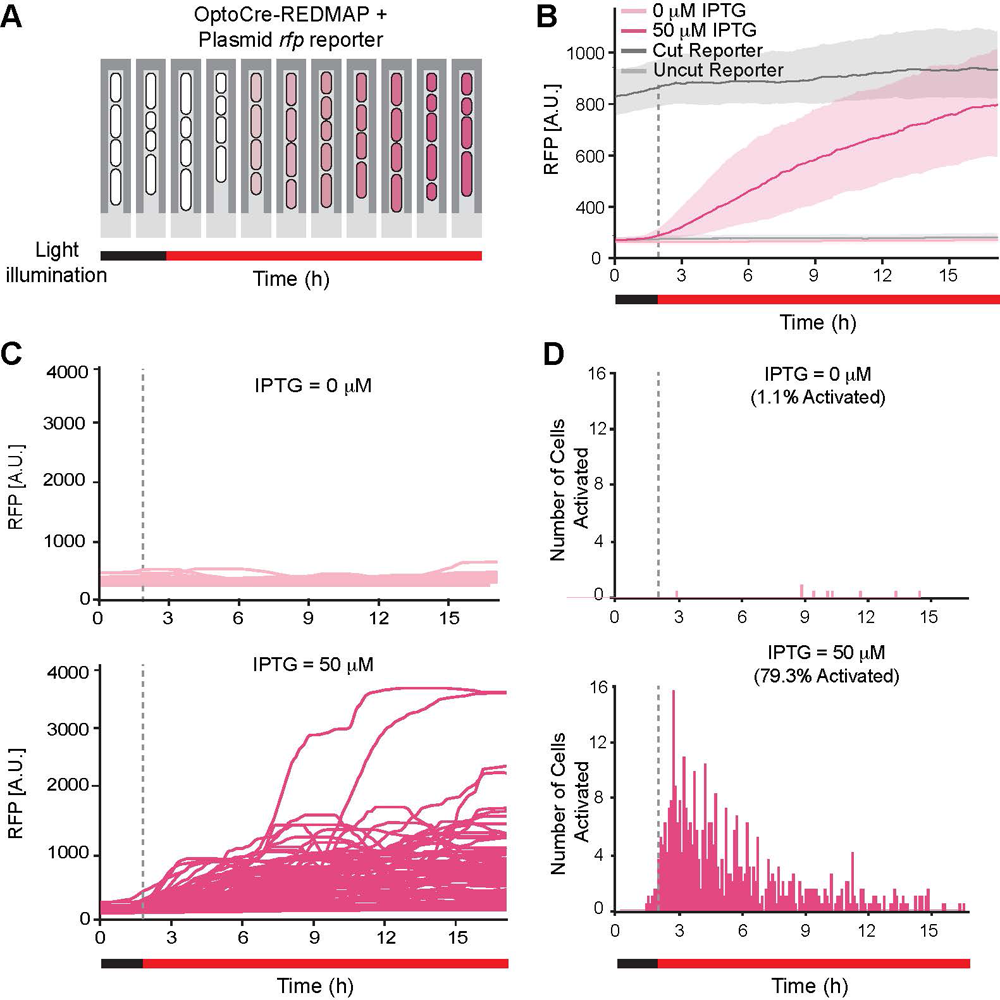
Activation of gene expression in single cells over time. **(A)** Schematic illustrating RFP induction in response to red light over time. Cells are grown in the mother machine microfluidic device and the “mother” cell at the top is tracked over many hours. **(B)** Mean fluorescence across all imaged mother cells. In all cases, cells were exposed to red light illumination starting at t = 3 h. The 0 and 50 µM IPTG conditions correspond to cells without and with OptoCre-REDMAP expression with the plasmid *rfp* reporter. Shaded error bars show standard deviation around the mean fluorescence over time. n ≥ 800 cells per condition. A.U., arbitrary units. **(C)** Fluorescence over time for individual cells with 0 and 50 µM IPTG induction. Data from the “mother” cells (i.e. the cells at the top of the chamber) are shown. **(D)** Time of single-cell RFP activation for individual cells with 0 and 50 µM IPTG induction.

We were able to successfully activate cells within the microfluidic device using this protocol, with the 0 µM IPTG condition remaining close to the uncut reporter negative control, while the 50 µM IPTG condition reached the levels of the cut reporter positive control (Fig. 4B). However, as we observed in the snapshot experiments, increases in fluorescence were slow to appear. Interestingly, single-cell data suggest that there is significant heterogeneity in the time of fluorescence expression activation following light exposure (Fig. 4C, Movies S1-S2). Notably in the 50 µM IPTG condition, the time to initiate expression varies widely between cells. To quantify activation time, we set a heuristic threshold corresponding to twice the median fluorescence of the uncut reporter, and recorded the time when individual cells exceeded this value. Activation time was highly variable and, in some cases, lagged light exposure by many hours (Fig. 4D). Ultimately, most cells were activated (79.3%) in the 50 µM IPTG case, which is notably higher than 1.1% activation in the 0 µM case. Single-cell growth rates were similar across all conditions and timepoints and showed no correlation with the time when RFP expression was activated (Fig. S9). We also asked whether confounding factors such as IPTG uptake rate could be causing this variation. To test this, we used a strain with an IPTG-inducible copy of *rfp*. Under these conditions we observed activation of all cells within 1-2 hours following induction with little variation between cells, indicating that the induction system has little to do with this variation (Fig. S10). These results, combined with the snapshot data, strongly suggest that light-induced gene expression by optogenetic Cre is heterogeneous across individual cells, with large variations in the timing of activation.

## Discussion

In this study, we systematically compared three bacterial optogenetic Cre systems using multiple reporters at both the population and single-cell levels. Each system showed distinct performance characteristics depending on the reporter used, with OptoCre-REDMAP and OptoCre-Vvd demonstrating efficient activation with the plasmid *rfp* reporter, while PA-Cre was most effective when paired with the chromosomal *mCherry-cat* reporter. All three recombinases performed well on a hybrid chromosomal *rfp* reporter. Our findings suggest that Cre variants may not be perfectly interchangeable, despite recognizing the same loxP sequence. Among the three optogenetic Cre systems we tested, OptoCre-REDMAP was the most versatile, demonstrating recombination when paired with all three reporters. However, it is worth emphasizing that the particular use case is likely to be critical for informing the selection of recombinase-reporter pairs, as the systems differ in what light wavelength activates them, the placement of the reporter, and whether expression is activated or genes are excised.

We anticipate that future studies will capitalize on the ability to use spatially-patterned light to target individual cells, thus we extended our population-level analysis to include single-cell results. These experiments revealed significant heterogeneity in recombination by optogenetic Cre, with substantial delays between light induction and reporter expression. In the case of the two chromosomal reporters, underlying fluorescence distributions were often bimodal, with cells shifting between the two states as a function of light exposure. qPCR analysis revealed that the recombination efficiency of OptoCre-REDMAP with the plasmid *rfp* reporter was 79.9% after 4 h of light exposure. This incomplete excision likely contributes to variability and delays in reporter activation. Split recombinases themselves may exhibit reduced functionality and stability, which could influence their enzymatic kinetics by altering both the timing and rate of reporter excision. In addition, we observed substantial temporal heterogeneity in system performance. Using OptoCre-REDMAP with the plasmid *rfp* reporter, we observed that the maximum fluorescence associated with low light exposure durations was not achieved until 18 hours post-illumination, despite some cells activating within minutes of red light exposure. Experiments using the mother machine device confirmed this delayed activation pattern and revealed substantial variability in activation timing following light exposure. This heterogeneity in activation dynamics has important implications for single-cell applications where precise temporal control is often desired.

There are several future directions for this work that could reveal additional sources of variability or improve single-cell precision. Generally, characterization and optimization efforts could focus on the optogenetic Cre components, the reporter, the methodology, or some combination of these factors. For example, it would be interesting to swap components within the optogenetic Cre constructs, such a switching photodimers, to better understand the constraints they introduce. Analysis of the single-cell data revealed examples of leaky activation in the dark state, and it would be interesting to test alternative genetic designs that reduce leakiness and increase the dynamic range of expression. While we assessed the single-cell activation dynamics of OptoCre-REDMAP using mother machine, extending this analysis to other optogenetic Cre systems could provide insight into the differences between the systems and how variation emerges. Also, in the reporter-only controls, we observed substantial cell-to-cell variation (e.g. for cut plasmid *rfp* reporter, Fig. 2A), indicating that optimizing reporter designs to reduce variation could mitigate some of the heterogeneity we encountered. Future experiments could investigate alternative causes for variance in both plasmid and chromosomal reporters, including factors such as light penetration, cell-cycle phase, metabolic states, Cre expression noise, and genetic construct design. There are also opportunities to optimize the experimental protocols, such as changing light intensities or the frequency of illumination^33,34,47^. The REDMAP system dissociates under far-red light, so controlling the ratio of red to far-red light is another direction for optimization.

It will also be interesting to explore differences between optogenetic Cre and non-optogenetic versions^48,49^. For example, testing how the distance between loxP sites impacts recombination, as we observed substantial differences in function for some of the optogenetic recombinases when switching between the chromosomal *mCherry-cat* and *rfp* reporters. Prior studies have characterized distance effects with non-optogenetic versions of Cre^50,51^ and it will be valuable to extend this to optogenetic designs.

OptoCre-REDMAP, OptoCre-Vvd, and PA-Cre are all powerful tools for bacterial optogenetics, and bulk culture assays have begun to demonstrate their potential. Our single-cell resolution results reveal key insights into the performance of these systems, suggesting the presence of bimodal distributions and substantial delays in expression of reporter genes that can vary across populations of cells. These results have critical implications as we move towards experiments that fully capitalize on the potential of dynamic, spatial-targeting with light for single-cell control.

## Materials and methods

### Strains and plasmids

All experiments use *E. coli* MG1655. The three recombinase constructs are all on plasmids with a low copy pSC101 origin of replication and an ampicillin resistance gene. In all cases, the recombinase genes are under the control of IPTG-inducible promoters: P_lacUV5_ for OptoCre-REDMAP and OptoCre-Vvd and P_LlacO1_ for PA-Cre, unless otherwise noted. All three optogenetic recombinases are variants on iCre^48^ and are split so that the nCre fragment consists of the first 43 amino acids of Cre, with cCre containing the remainder. The OptoCre-REDMAP plasmid was derived from Jafarbeglou et al.^33^. In the original study, the *ho1-pcy* gene encoding PCB was included on the reporter plasmid. For this study, we moved *ho1-pcy* to the plasmid containing OptoCre-REDMAP (Fig. S1) so that we could use the same reporters for all three recombinases. We compared the activation of both versions of the OptoCre-REDMAP system and confirmed that the fold-change and growth were similar, regardless of the placement of *ho1-pcy* (Fig. 1C, Fig. S11). The OptoCre-Vvd plasmid was sourced from Sheets et al.^34^ (AddGene #160400). PA-Cre was a gift from Dr. Yuichi Wakamoto^18^. To test the effects of promoter and RBS strength on the functionality of the optogenetic Cre system, we also cloned PA-Cre into the same backbone as OptoCre-Vvd and OptoCre-REDMAP. In cases where we used a non-optogenetic Cre, the construct is a non-split version of iCre controlled by the P_lacUV5_ promoter. In experiments requiring a non-functional Cre (dCre), we used iCreK201A under the control of the P_lacUV5_ promoter, derived from Gosh et al.^44^.

The plasmid *rfp* reporter is from Sheets et al.^34^ (AddGene #134405) and contains a medium copy p15A origin of replication with kanamycin resistance gene with *mrfp1*, which we refer to as *rfp* throughout the manuscript. The chromosomal *mcherry-cat* reporter was a gift from Dr. Yuichi Wakamoto^18^. We created the chromosomal *rfp* reporter using chromosomal integration, where the reporter construct and a kanamycin nucleotidyltransferase (*knt*) cassette, flanked by FRT sites, were inserted downstream of *nupG* using the lambda red recombinase system^52^. Kanamycin selection was used to identify cells with successful integration, then the cassette was cured using a plasmid encoding FLP recombinase with a temperature sensitive origin of replication following the protocol from Datsenko and Wanner^52^. The temperature sensitive plasmid was then cured prior to experiments. For both the plasmid and chromosomal *rfp* reporters, *rfp* is under the control of a P_W4_ constitutive promoter as described in Sheets et al.^34^.

The uncut reporter control consists of cells with the reporter only and no recombinase plasmid. The cut reporter control contains the reporter with a single loxP site and no recombinase plasmid. In the case of the plasmid *rfp* reporter we created two versions: one where we cloned a single loxP site between the constitutive promoter and *rfp* (denoted ‘cut reporter’) and a second where we used a non-optogenetic version of Cre to excise the loxP flanked terminator (denoted ‘Cre cut reporter’). For the chromosomal *mCherry-cat* and *rfp* reporters, we used non-optogenetic Cre to generate the cut versions of the reporters. To test direct IPTG induction of fluorophore expression in the mother machine, we used MG1655 cells with an IPTG-inducible reporter plasmid encoding RFP under the control of the P_lacUV5_ promoter from Lee et al.^53^.

### Light exposure assay

All liquid culture experiments were conducted at 37°C with 220 rpm shaking. Cultures were maintained in the dark except during designated light exposure periods. Strains were grown overnight from a single colony in LB medium with 100 μg/mL carbenicillin and 30 μg/mL kanamycin for plasmid maintenance. The next day, cultures were refreshed 1:100 in M9 medium (M9 salts supplemented with 2 mM MgSO_4_, 0.4% glycerol, 0.2% casamino acids, 100 µM CaCl_2_) with 100 μg/mL carbenicillin and 30 μg/mL kanamycin and induced for 2 h with 100 μM IPTG. Light exposure was performed using an OptoPlate-96 device equipped with LEDs emitting light at a wavelength of 630 nm (red) or 465 nm (blue) with an intensity of 155 μW/cm^2^ per well. The OptoPlate-96 was built following the design by Bugaj et al.^54^ using the protocol by Dunlop^55^.

After a 2 h IPTG induction period for the experiments in Fig. 1 and 2, cultures were kept in the dark until the designated exposure time, after which they were subjected to either red (OptoCre-REDMAP) or blue (OptoCre-Vvd, PA-Cre) light. After light exposure, cultures were transferred to M9 medium with 100 μg/mL carbenicillin and 30 μg/mL kanamycin for plasmid maintenance and grown overnight (∼18 h). For population-level experiments, plate reader data were collected using a BioTek Synergy H1 device. Optical density absorbance readings were collected at 600 nm (OD600). Fluorescence readings use an excitation of 584 nm and emission of 610 nm. Single-cell snapshot protocols are described below.

### Antibiotic susceptibility assays

Antibiotic susceptibility assays were conducted using the protocol in Sheets et al.^35^. Chloramphenicol antibiotic stocks were prepared by dissolving the antibiotic in 99% ethanol, with concentrations normalized for potency based on CLSI standards^56^. Assay plates for antibiotic resistance measurement were prepared by serially diluting chloramphenicol in 150 μL M9 medium in 96 well plates. After light exposure, cells were diluted 1:30 into the 96 well plates in triplicate then grown overnight. OD600 measurements were collected using a BioTek Synergy H1 plate reader.

### Microscopy and image analysis

After light illumination, cells were refreshed in M9 medium with 100 μg/mL carbenicillin and 30 μg/mL kanamycin and grown overnight. The following day, cells were refreshed 1:100 in M9 medium with antibiotics for plasmid maintenance for 2 h. Cells were then placed on 1.5% low melting agarose pads made with M9 medium. We conducted single-cell imaging using a Nikon Ti2-E microscope with a 100x objective. Images were segmented and analyzed using DeLTA^57^. For the experiments in Fig. 3, we took microscopy images at intermediate timepoints, including 0, 1, and 3 h after light exposure in addition to the overnight (18 h) timepoint. For the 3 h timepoint, we diluted cells 1:2 in M9 medium prior to imaging to avoid overcrowding of the field of view.

### Colony PCR and qPCR

For colony PCR, after light illumination cells were serially diluted 1:30 four times and plated onto agar containing the appropriate maintenance antibiotics for the reporter (30 μg/mL kanamycin for plasmid reporter and no antibiotics for the chromosomal reporters). Plates were incubated at 37°C overnight to allow single cells to develop into colonies. Five random colonies were picked from the 0 h and 4 h light conditions for colony PCR using the DreamTaq polymerase (Thermo Fisher Scientific, Waltham, MA, USA). Primers are listed in Table S1. PCR products were run on a 1% agarose gel.

For qPCR, we generated a standard curve with known amounts of the uncut reporter plasmid, ranging from 100 ng to 1 pg. After light illumination, cells were refreshed 1:100 in 3 mL of M9 medium with both 100 μg/mL carbenicillin and 30 μg/mL kanamycin. After overnight growth, the plasmid DNA was purified and collected using the GenCatch Plasmid DNA Mini-prep kit (Epoch Life Science Inc., Missouri City, TX, USA). The resulting DNA was diluted to 10 ng/μL and a volume of 0.5 μL of the diluted DNA was used for the qPCR reaction with a KAPA SYBR FAST Universal qPCR kit (Roche, Basel, Switzerland). For each condition, we included technical three replicates. Primers used for qPCR are listed in Table S1.

### Mother machine microfluidic device

The mother machine device is based on the design used in Sampaio et al.^12^. SU-resin molds were made via stereolithography based on the design available at https://gitlab.com/dunloplab/mother_machine. Polydimethylsiloxane (PDMS) (Dow Corning Sylgard 184) was poured into the mold and cured overnight at 75 °C. Individual chips were cut out and the inlet and outlet holes were cut using a 0.75 mm biopsy punch. The chip was plasma bonded to a glass slide and incubated at 75 °C overnight before use.

Overnight cell cultures were refreshed 1:100 in LB medium with 0.2 g/L Pluronic F-127 to reduce cell adhesion to the device. After 2-3 hours of growth, cells were loaded into the mother machine device by centrifugation at 5000 rpm for 3 min. Media tubing was connected at the inlet and outlet ports and LB was pumped to the chip using a peristaltic pump at a rate of 20 μL/min. 50 μg/mL carbenicillin and 15 μg/mL kanamycin were added to the media for plasmid maintenance. 50 μM of IPTG was used for induction. After 3 hours of recovery, we began imaging using a Nikon Ti2-E microscope with 100x objective. After 2 hours of imaging, we started red light illumination, which was continued through the end of the experiment. Red light was provided by a LED ring light (NeoPixel Ring - 24 x 5050 RGBW LEDs) placed above the microscope stage. For red light illumination we used a 60 sec period of light exposure for each 5 min imaging period. Images were segmented and analyzed using DeLTA^57^. We calculated the time of activation by selecting a heuristic threshold of 362 A.U., which corresponds to twice the median fluorescence of the uncut reporter. After the fluorescence of a cell exceeded this threshold and stayed above the threshold for at least 6 frames (30 min) it was categorized as activated.

## Acknowledgements

We thank Dr. Yuichi Wakamoto for kindly sharing the PA-Cre system and chromosomal *mCherry-cat* reporter. Thanks to Cristian Coriano-Ortiz for helpful discussions about quantifying recombination efficiencies. This work was supported by NIH Grant R01AI102922, NSF Grant MCB-2324909, and a Kilachand Award made through the Multicellular Design Program at Boston University.

## References

1. Zhao, E.M., Zhang, Y., Mehl, J., Park, H., Lalwani, M.A., Toettcher, J.E., and Avalos, J.L. (2018). Optogenetic regulation of engineered cellular metabolism for microbial chemical production. Nature 555, 683–687. 10.1038/nature26141.

2. Pu, L., Yang, S., Xia, A., and Jin, F. (2018). Optogenetics Manipulation Enables Prevention of Biofilm Formation of Engineered Pseudomonas aeruginosa on Surfaces. ACS Synth. Biol. 7, 200–208. 10.1021/acssynbio.7b00273.

3. Milias-Argeitis, A., Rullan, M., Aoki, S.K., Buchmann, P., and Khammash, M. (2016). Automated optogenetic feedback control for precise and robust regulation of gene expression and cell growth. Nat. Commun. 7, 12546. 10.1038/ncomms12546.

4. Zhao, E.M., Suek, N., Wilson, M.Z., Dine, E., Pannucci, N.L., Gitai, Z., Avalos, J.L., and Toettcher, J.E. (2019). Light-based control of metabolic flux through assembly of synthetic organelles. Nat. Chem. Biol. 15, 589–597. 10.1038/s41589-019-0284-8.

5. Chia, N., Lee, S.Y., and Tong, Y. (2022). Optogenetic tools for microbial synthetic biology. Biotechnol. Adv. 59, 107953. 10.1016/j.biotechadv.2022.107953.

6. van Gestel, J., Bareia, T., Tenennbaum, B., Dal Co, A., Guler, P., Aframian, N., Puyesky, S., Grinberg, I., D’Souza, G.G., Erez, Z., et al. (2021). Short-range quorum sensing controls horizontal gene transfer at micron scale in bacterial communities. Nat. Commun. 12, 2324. 10.1038/s41467-021-22649-4.

7. Van Vliet, S., Hauert, C., Fridberg, K., Ackermann, M., and Co, A.D. (2022). Global dynamics of microbial communities emerge from local interaction rules. PLOS Comput. Biol. 18, e1009877. 10.1371/JOURNAL.PCBI.1009877.

8. Le Bec, M., Pouzet, S., Cordier, C., Barral, S., Scolari, V., Sorre, B., Banderas, A., and Hersen, P. (2024). Optogenetic spatial patterning of cooperation in yeast populations. Nat. Commun. 15, 75. 10.1038/s41467-023-44379-5.

9. Banderas, A., Hofmann, M., Cordier, C., Le Bec, M., Elizondo-Cantú, M.C., Chiron, L., Pouzet, S., Lifschytz, Y., Ji, W., Amir, A., et al. (2024). Optogenetic control of pheromone gradients and mating behavior in budding yeast. Preprint at Cold Spring Harbor Laboratory, 10.1101/2024.02.06.578657 https://doi.org/10.1101/2024.02.06.578657.

10. Quispe Haro, J.J., Chen, F., Los, R., Shi, S., Sun, W., Chen, Y., Idema, T., and Wegner, S.V. (2024). Optogenetic Control of Bacterial Cell-Cell Adhesion Dynamics: Unraveling the Influence on Biofilm Architecture and Functionality. Adv. Sci. 11, 2310079. 10.1002/advs.202310079.

11. Jin, X., and Riedel-Kruse, I.H. (2024). Optogenetic patterning generates multi-strain biofilms with spatially distributed antibiotic resistance. Nat. Commun. 15, 9443. 10.1038/s41467-024-53546-1.

12. Sampaio, N.M.V., Blassick, C.M., Andreani, V., Lugagne, J.-B., and Dunlop, M.J. (2022). Dynamic gene expression and growth underlie cell-to-cell heterogeneity in Escherichia coli stress response. Proc. Natl. Acad. Sci. 119, e2115032119. 10.1073/pnas.2115032119.

13. Uphoff, S., Lord, N.D., Okumus, B., Potvin-Trottier, L., Sherratt, D.J., and Paulsson, J. (2016). Stochastic activation of a DNA damage response causes cell-to-cell mutation rate variation. Science 351, 1094–1097. 10.1126/science.aac9786.

14. Sweeney, K., and McClean, M.N. (2023). Transcription factor localization dynamics and DNA binding drive distinct promoter interpretations. Cell Rep. 42, 112426. 10.1016/j.celrep.2023.112426.

15. Lugagne, J.-B., and Dunlop, M.J. (2019). Cell-machine interfaces for characterizing gene regulatory network dynamics. Curr. Opin. Syst. Biol. 14, 1–8. 10.1016/j.coisb.2019.01.001.

16. Scott, T.D., Sweeney, K., and McClean, M.N. (2019). Biological signal generators: integrating synthetic biology tools and in silico control. Curr. Opin. Syst. Biol. 14, 58–65. 10.1016/j.coisb.2019.02.007.

17. Fox, Z.R., Fletcher, S., Fraisse, A., Aditya, C., Sosa-Carrillo, S., Petit, J., Gilles, S., Bertaux, F., Ruess, J., and Batt, G. (2022). Enabling reactive microscopy with MicroMator. Nat. Commun. 2022 131 13, 1–8. 10.1038/s41467-022-29888-z.

18. Koganezawa, Y., Umetani, M., Sato, M., and Wakamoto, Y. (2022). History-dependent physiological adaptation to lethal genetic modification under antibiotic exposure. eLife 11. 10.7554/eLife.74486.

19. Lugagne, J.-B., Blassick, C.M., and Dunlop, M.J. (2024). Deep model predictive control of gene expression in thousands of single cells. Nat. Commun. 15, 2148. 10.1038/s41467-024-46361-1.

20. Chait, R., Ruess, J., Bergmiller, T., Tkačik, G., and Guet, C.C. (2017). Shaping bacterial population behavior through computer-interfaced control of individual cells. Nat. Commun. 8, 1535. 10.1038/s41467-017-01683-1.

21. McQuillen, R., Perez, A.J., Yang, X., Bohrer, C.H., Smith, E.L., Chareyre, S., Tsui, H.-C.T., Bruce, K.E., Hla, Y.M., McCausland, J.W., et al. (2024). Light-dependent modulation of protein localization and function in living bacteria cells. Nat. Commun. 15, 10746. 10.1038/s41467-024-54974-9.

22. Cannarsa, M.C., Liguori, F., Pellicciotta, N., Frangipane, G., and Leonardo, R.D. (2024). Light-driven synchronization of optogenetic clocks. eLife 13. 10.7554/eLife.97754.2.

23. Goglia, A.G., and Toettcher, J.E. (2019). A bright future: optogenetics to dissect the spatiotemporal control of cell behavior. Curr. Opin. Chem. Biol. 48, 106–113. 10.1016/j.cbpa.2018.11.010.

24. Hoffman, S.M., Tang, A.Y., and Avalos, J.L. (2022). Optogenetics Illuminates Applications in Microbial Engineering. Annu. Rev. Chem. Biomol. Eng. 13, 373–403. 10.1146/annurev-chembioeng-092120-092340.

25. Hoess, R.H., and Abremski, K. (1985). Mechanism of strand cleavage and exchange in the Cre-*lox* site-specific recombination system. J. Mol. Biol. 181, 351–362. 10.1016/0022-2836(85)90224-4.

26. Weinberg, B.H., Cho, J.H., Agarwal, Y., Pham, N.T.H., Caraballo, L.D., Walkosz, M., Ortega, C., Trexler, M., Tague, N., Law, B., et al. (2019). High-performance chemical- and light-inducible recombinases in mammalian cells and mice. Nat. Commun. 10, 4845. 10.1038/s41467-019-12800-7.

27. Tous, C., Kinstlinger, I.S., Rice, M.E.L., Deng, J., and Wong, W.W. (2025). Multiplexing light-inducible recombinases to control cell fate, Boolean logic, and cell patterning in mammalian cells. Sci. Adv. 11, eadt1971. 10.1126/sciadv.adt1971.

28. Short, A.E., Kim, D., Milner, P.T., and Wilson, C.J. (2023). Next generation synthetic memory via intercepting recombinase function. Nat. Commun. 14, 5255. 10.1038/s41467-023-41043-w.

29. Meador, K., Wysoczynski, C.L., Norris, A.J., Aoto, J., Bruchas, M.R., and Tucker, C.L. (2019). Achieving tight control of a photoactivatable Cre recombinase gene switch: new design strategies and functional characterization in mammalian cells and rodent. Nucleic Acids Res. 47, e97. 10.1093/nar/gkz585.

30. Zhou, Y., Kong, D., Wang, X., Yu, G., Wu, X., Guan, N., Weber, W., and Ye, H. (2022). A small and highly sensitive red/far-red optogenetic switch for applications in mammals. Nat. Biotechnol. 40, 262–272. 10.1038/s41587-021-01036-w.

31. L-SCRaMbLE as a tool for light-controlled Cre-mediated recombination in yeast | Nature Communications https://www.nature.com/articles/s41467-017-02208-6.

32. Dufour, A., Duplus-Bottin, H., Boukéké-Lesplulier, T., Casassa, E., Triqueneaux, G., Darthenay-Kiennemann, C., Dumont, A., Moali, C., Vittoz, F., Jost, D., et al. (2024). Kinetic properties of optogenetic DNA editing by LiCre-loxP. Preprint at bioRxiv, 10.1101/2024.05.17.594525

33. Jafarbeglou, F., and Dunlop, M.J. (2024). Red Light Responsive Cre Recombinase for Bacterial Optogenetics. ACS Synth. Biol. 13, 3991–4001. 10.1021/acssynbio.4c00388.

34. Sheets, M.B., Wong, W.W., and Dunlop, M.J. (2020). Light-Inducible Recombinases for Bacterial Optogenetics. ACS Synth. Biol. 9, 227–235. 10.1021/acssynbio.9b00395.

35. Sheets, M.B., Tague, N., and Dunlop, M.J. (2023). An optogenetic toolkit for light-inducible antibiotic resistance. Nat. Commun. 14, 1034. 10.1038/s41467-023-36670-2.

36. Lim, C.K., Yeoh, J.W., Kunartama, A.A., Yew, W.S., and Poh, C.L. (2023). A biological camera that captures and stores images directly into DNA. Nat. Commun. 14, 3921. 10.1038/s41467-023-38876-w.

37. Sharrock, R.A., and Quail, P.H. (1989). Novel phytochrome sequences in Arabidopsis thaliana: structure, evolution, and differential expression of a plant regulatory photoreceptor family. Genes Dev. 3, 1745–1757. 10.1101/gad.3.11.1745.

38. Loros, J.J., and Dunlap, J.C. (2001). Genetic and Molecular Analysis of Circadian Rhythms in Neurospora. Annu. Rev. Physiol. 63, 757–794. 10.1146/annurev.physiol.63.1.757.

39. Zoltowski, B.D., Schwerdtfeger, C., Widom, J., Loros, J.J., Bilwes, A.M., Dunlap, J.C., and Crane, B.R. (2007). Conformational Switching in the Fungal Light Sensor Vivid. Science 316, 1054–1057. 10.1126/science.1137128.

40. Kawano, F., Okazaki, R., Yazawa, M., and Sato, M. (2016). A photoactivatable Cre–loxP recombination system for optogenetic genome engineering. Nat. Chem. Biol. 12, 1059–1064. 10.1038/nchembio.2205.

41. Kawano, F., Suzuki, H., Furuya, A., and Sato, M. (2015). Engineered pairs of distinct photoswitches for optogenetic control of cellular proteins. Nat. Commun. 6, 6256. 10.1038/ncomms7256.

42. Malan, T.P., and McClure, W.R. (1984). Dual promoter control of the escherichia coli lactose operon. Cell 39, 173–180. 10.1016/0092-8674(84)90203-4.

43. Shaw, W.V. (1967). The Enzymatic Acetylation of Chloramphenicol by Extracts of R Factor-resistant *Escherichia coli*. J. Biol. Chem. 242, 687–693. 10.1016/S0021-9258(18)96259-9.

44. Ghosh, K., Guo, F., and Van Duyne, G.D. (2007). Synapsis of *loxP* Sites by Cre Recombinase*. J. Biol. Chem. 282, 24004–24016. 10.1074/jbc.M703283200.

45. Balleza, E., Kim, J.M., and Cluzel, P. (2018). Systematic characterization of maturation time of fluorescent proteins in living cells. Nat. Methods 15, 47–51. 10.1038/nmeth.4509.

46. Wang, P., Robert, L., Pelletier, J., Dang, W.L., Taddei, F., Wright, A., and Jun, S. (2010). Robust growth of escherichia coli. Curr. Biol. 20, 1099–1103. 10.1016/j.cub.2010.04.045.

47. Lee, J.B., Caywood, L.M., Lo, J.Y., Levering, N., and Keung, A.J. (2021). Mapping the dynamic transfer functions of eukaryotic gene regulation. Cell Syst. 12, 1079–1093.e6. 10.1016/j.cels.2021.08.003.

48. Shimshek, D. r., Kim, J., Hübner, M. r., Spergel, D. j., Buchholz, F., Casanova, E., Stewart, A. f., Seeburg, P. h., and Sprengel, R. (2002). Codon-improved Cre recombinase (iCre) expression in the mouse. genesis 32, 19–26. 10.1002/gene.10023.

49. Garabello, E., Yoon, H., Reid, M.C., and Giometto, A. (2024). Tunable Low-Rate Genomic Recombination with Cre-lox in E. coli: A Versatile Tool for Synthetic Biology and Environmental Sensing. Preprint at bioRxiv, 10.1101/2024.10.02.616356 https://doi.org/10.1101/2024.10.02.616356.

50. Hoess, R., Wierzbicki, A., and Abremski, K. (1985). Formation of small circular DNA molecules via an in vitro site-specific recombination system. Gene 40, 325–329. 10.1016/0378-1119(85)90056-3.

51. Hosur, V., Erhardt, V., Hartig, E., Lorenzo, K., Megathlin, H., and Tarchini, B. (2024). Large-Scale Genome-Wide Optimization and Prediction of the Cre Recombinase System for Precise Genome Manipulation in Mice. Preprint at Research Square, 10.21203/rs.3.rs-4595968/v1 https://doi.org/10.21203/rs.3.rs-4595968/v1.

52. Datsenko, K.A., and Wanner, B.L. (2000). One-step inactivation of chromosomal genes in Escherichia coli K-12 using PCR products. Proc. Natl. Acad. Sci. 97, 6640–6645. 10.1073/pnas.120163297.

53. Lee, T.S., Krupa, R.A., Zhang, F., Hajimorad, M., Holtz, W.J., Prasad, N., Lee, S.K., and Keasling, J.D. (2011). BglBrick vectors and datasheets: A synthetic biology platform for gene expression. J. Biol. Eng. 5, 12. 10.1186/1754-1611-5-12.

54. Bugaj, L.J., and Lim, W.A. (2019). High-throughput multicolor optogenetics in microwell plates. Nat. Protoc. 14, 2205–2228. 10.1038/s41596-019-0178-y.

55. Dunlop, M.J. (2021). A Supplemental Guide to Building the optoPlate-96.

56. Wiegand, I., Hilpert, K., and Hancock, R.E.W. (2008). Agar and broth dilution methods to determine the minimal inhibitory concentration (MIC) of antimicrobial substances. Nat. Protoc. 2008 32 3, 163–175. 10.1038/nprot.2007.521.

57. O’Connor, O.M., Alnahhas, R.N., Lugagne, J.B., and Dunlop, M.J. (2022). DeLTA 2.0: A deep learning pipeline for quantifying single-cell spatial and temporal dynamics. PLOS Comput. Biol. 18, e1009797. 10.1371/JOURNAL.PCBI.1009797.

