## Supplementary Information for "Single-cell characterization of bacterial optogenetic Cre recombinases"

1. Supplementary Figures
2. Supplementary Tables
3. Supplementary Movie Captions

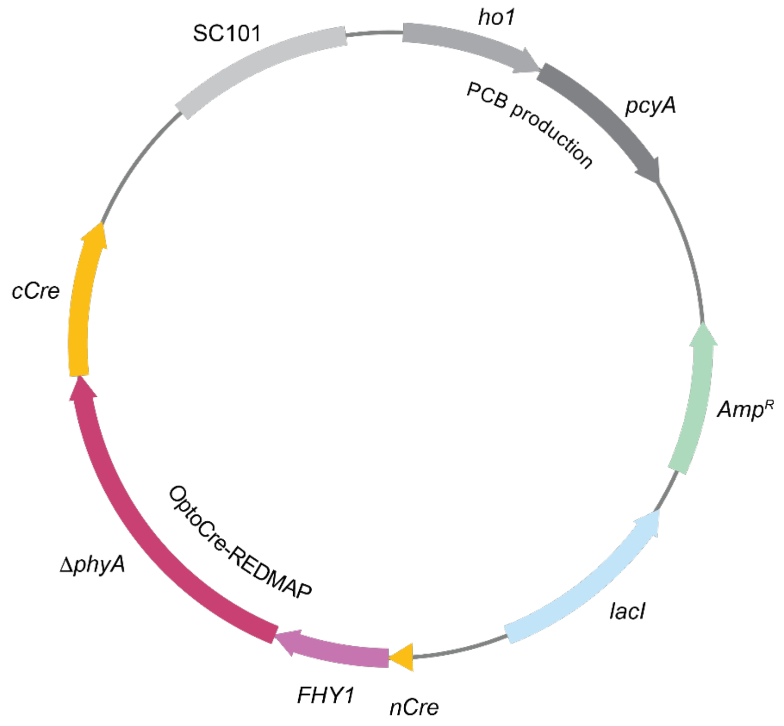


**Figure S1.** OptoCre-REDMAP with the CTG start codon. This plasmid also contains the *ho1-pcyA* gene that produces the chromophore phycocyanobilin (PCB).

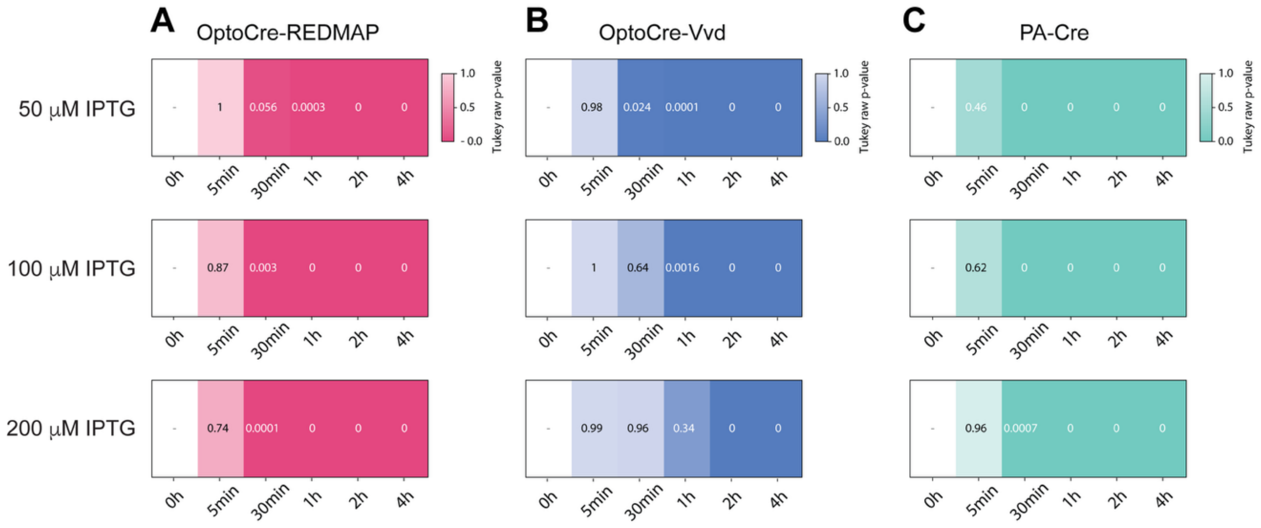


**Figure S2. Statistical analysis for all IPTG concentrations.** One-way ANOVA followed by Tukey's post-hoc test comparing the 0 h condition versus the other timepoints at different IPTG concentrations (50, 100, 200 µM) for the plasmid *rfp* reporter with **(A)** OptoCre-REDMAP, **(B)** OptoCre-Vvd, and **(C)** PA-Cre. Numbers indicated in the plot correspond to *p*-values.

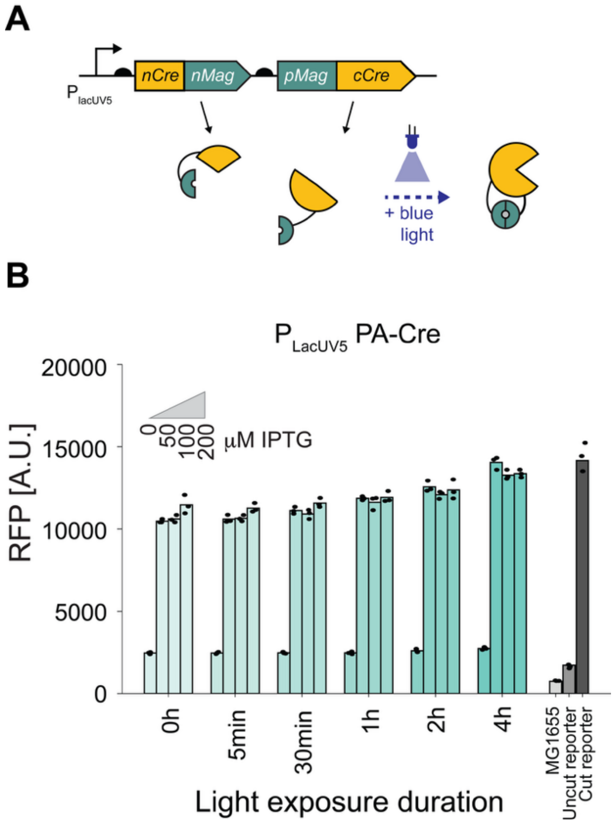


**Figure S3. (A)** The PA-Cre construct under the control of $P_{lacUV5}$. **(B)** RFP output of this construct under different light exposure conditions and IPTG concentrations. Bars represent the mean values, with individual biological replicates shown as dots (n = 3).

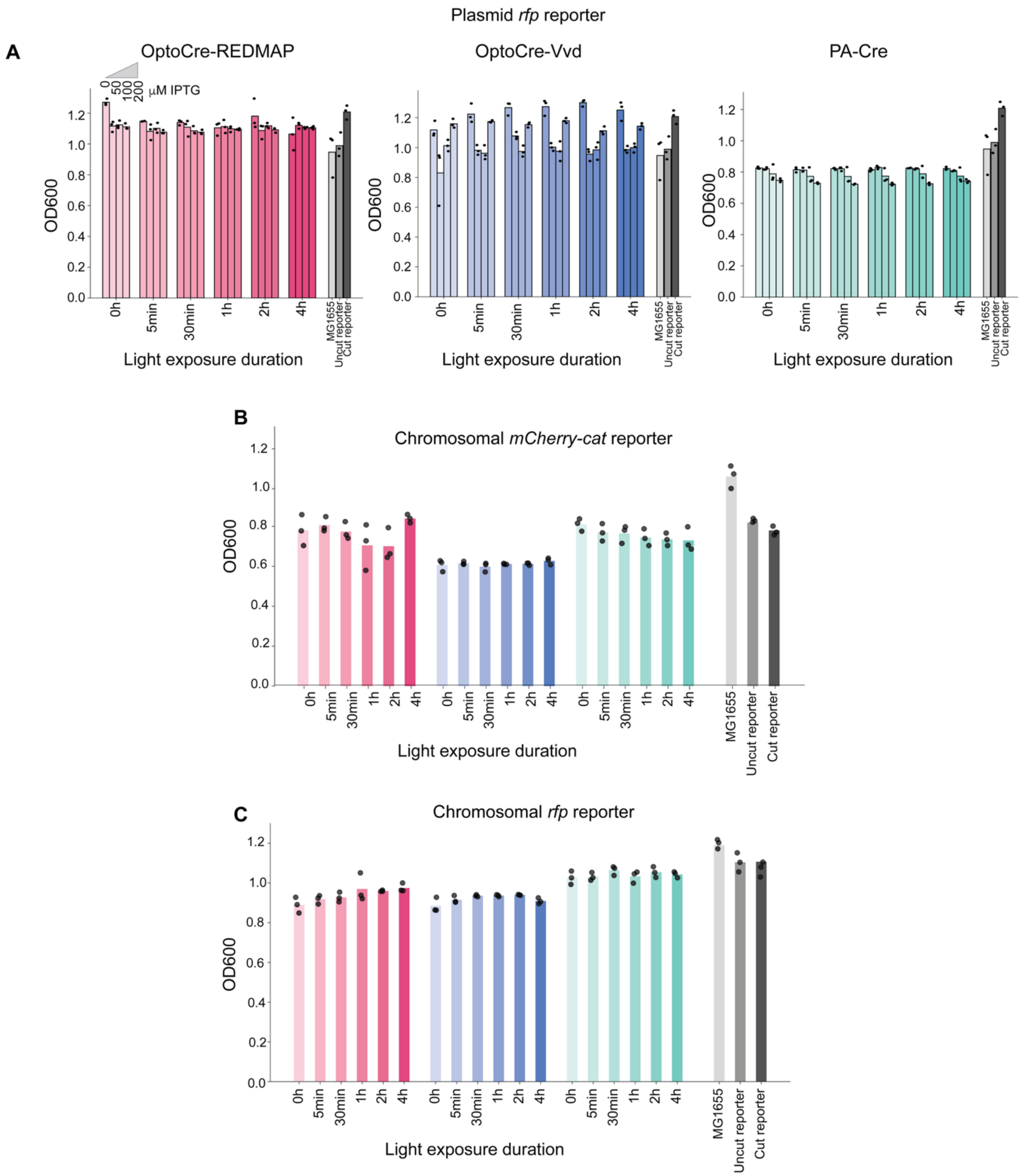


**Figure S4.** Optical density measured at 600 nm (OD600) for the three optogenetic Cre systems with different reporters after overnight growth. **(A)** Plasmid *rfp* reporter. Cells were induced with a range of IPTG concentrations (0, 50, 100, 200 µM). **(B)** Chromosomal *mCherry-cat* reporter. Cells were induced with 100 µM IPTG. **(C)** Chromosomal *rfp* reporter. Cells were induced with 100 µM IPTG. Bars represent the mean values, with individual biological replicates shown as dots (n = 3).

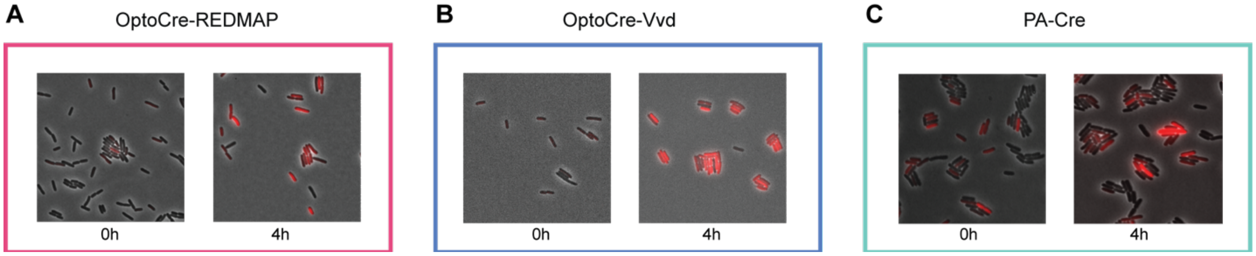


**Figure S5.** Representative microscopy snapshot images of the three optogenetic Cre systems. **(A)** OptoCre-REDMAP, **(B)** OptoCre-Vvd, and **(C)** PA-Cre with the plasmid *rfp* reporter after 0 or 4 h of light exposure.

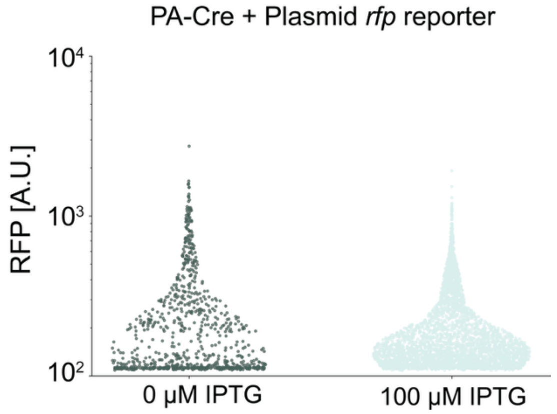


**Figure S6.** Single-cell RFP for PA-Cre with the plasmid *rfp* reporter without light exposure, with or without 100 µM IPTG.

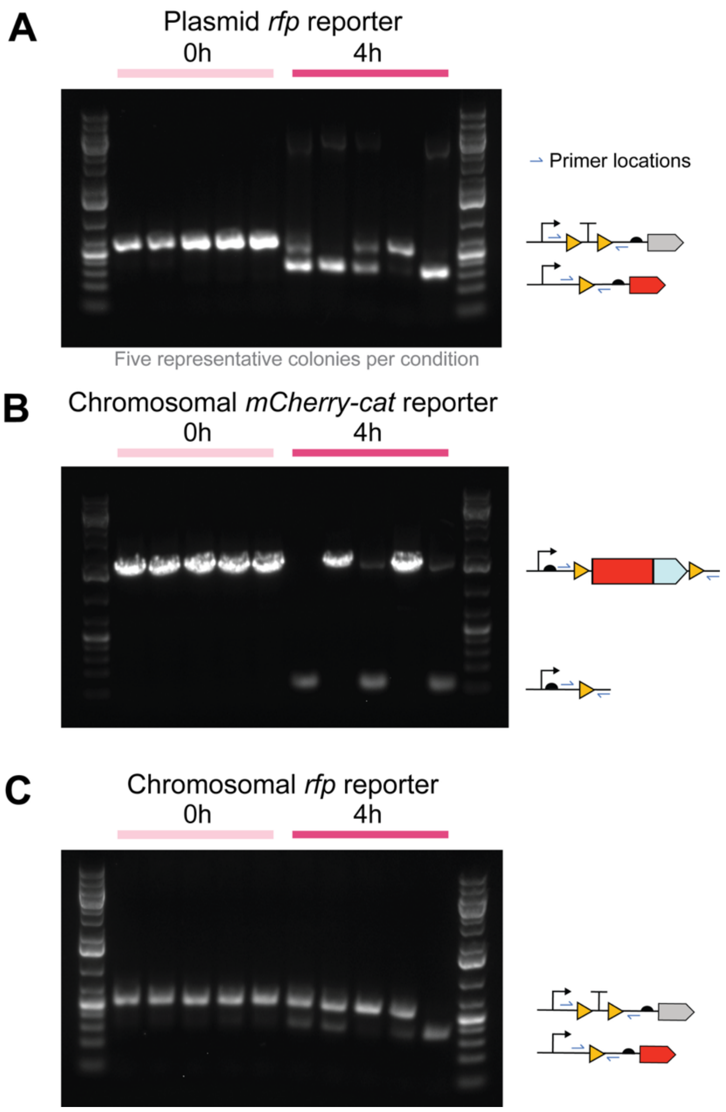


**Figure S7.** Colony PCR of five representative colonies each derived from single cells for OptoCre-REDMAP with 0 or 4 h of light exposure with all reporter types: **(A)** plasmid *rfp* reporter, **(B)** chromosomal *mCherry-cat* reporter, and **(C)** chromosomal *rfp* reporter. Schematics next to the gel images indicate uncut and cut versions of the reporter as well as primer locations for colony PCR.

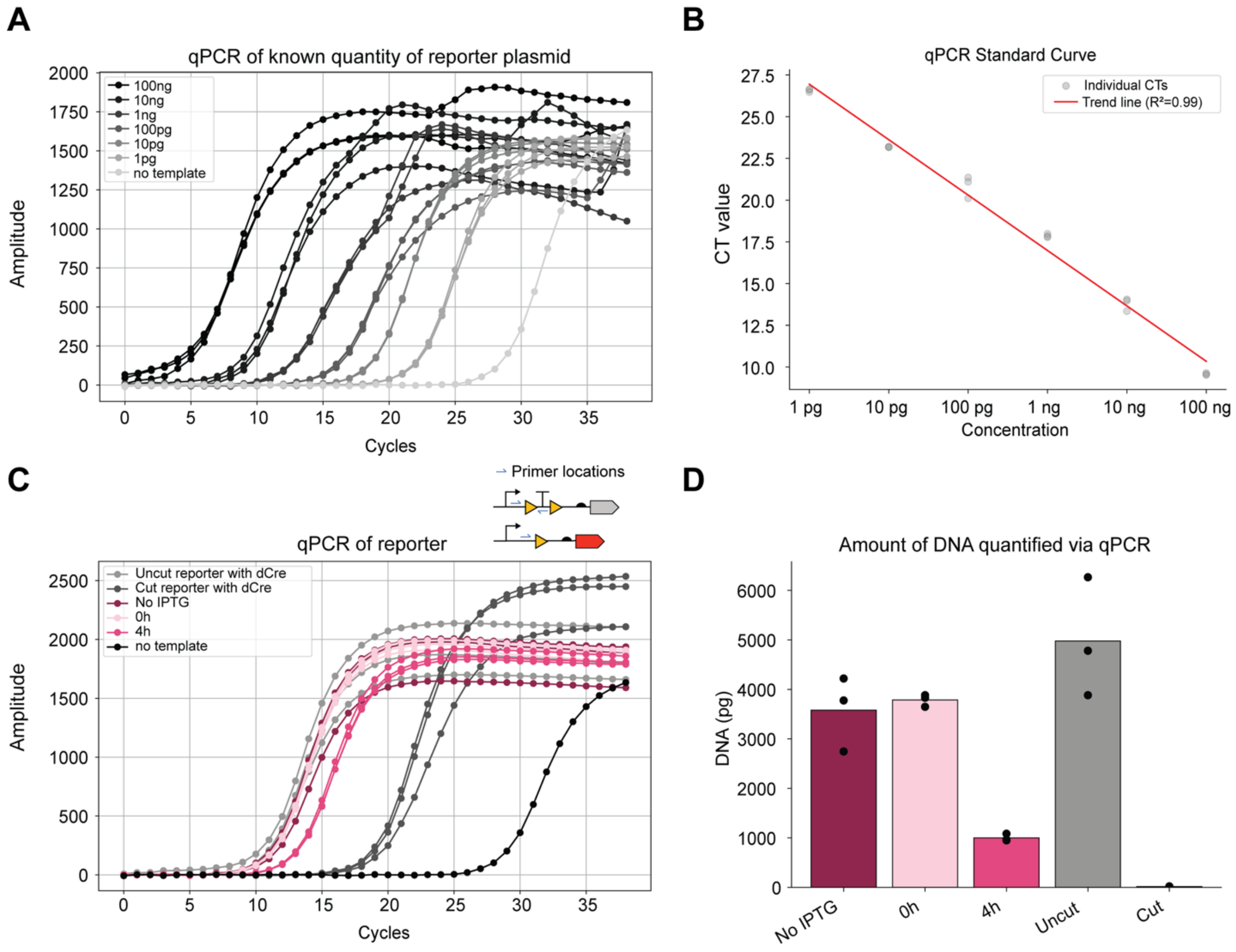


**Figure S8. qPCR quantification of recombination. (A)** qPCR of known amounts of the uncut plasmid *rfp* reporter. **(B)** Data from (A) used to generate a standard curve. **(C)** qPCR data of the plasmid *rfp* reporter after 0 h with no IPTG, 0 h with 100 µM IPTG, or 4 h of light exposure with 100 µM IPTG to determine the recombination efficiency of OptoCre-REDMAP. **(D)** Quantification of DNA based on the standard curve and qPCR results. Bars represent the mean values, with individual technical replicates shown as dots (n = 3).

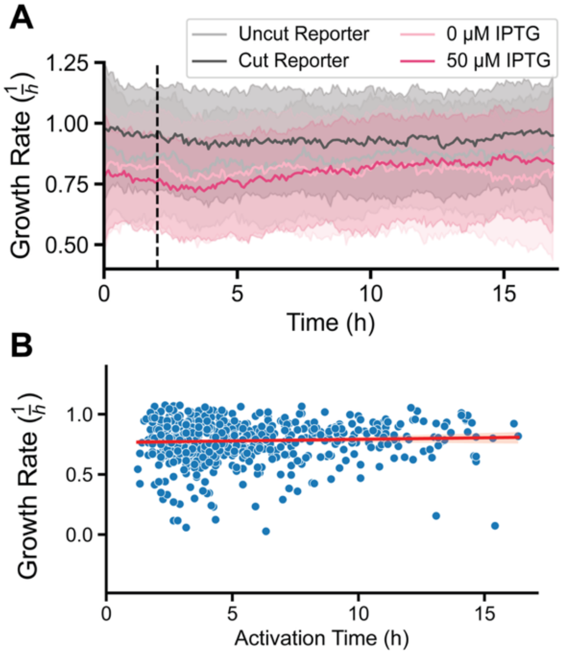


**Figure S9. (A)** Growth rate of single cells in the mother machine device over time. **(B)** Growth rate as a function of activation time. Pearson correlation analysis between activation time growth rate indicated no statistically significant linear relationship between the two parameters (r = 0.05 and $p$ = 0.19). Linear regression line is shown in red.

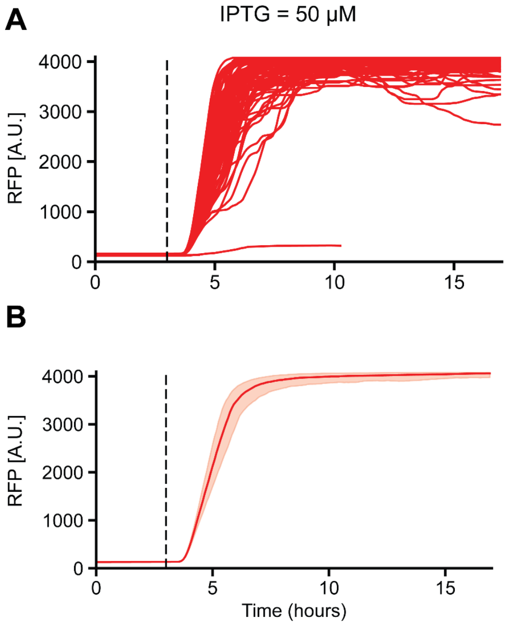


**Figure S10. (A)** Fluorescence over time for individual cells containing an IPTG inducible RFP on a plasmid ($P_{lacUV5}$-*rfp*). Data from the "mother" cells (i.e., the cells at the top of the chamber) are shown. 50 µM IPTG was added to the media starting at t = 3 h. **(B)** Mean fluorescence across all imaged cells.

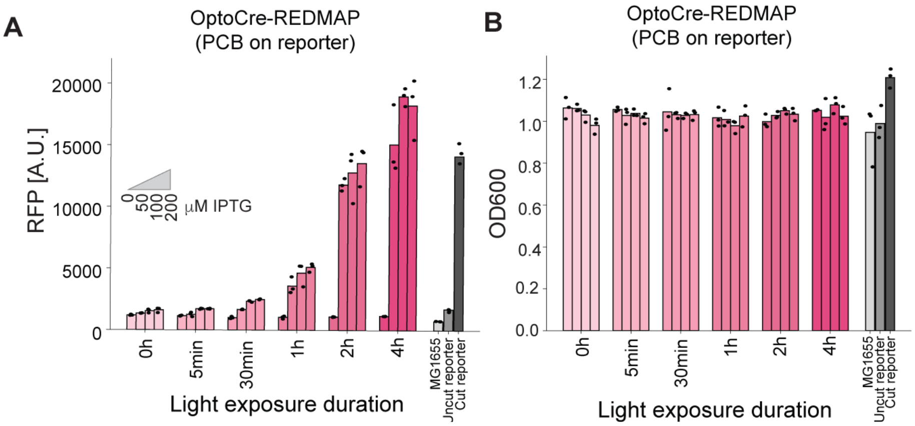


**Figure S11.** Population-level measurements of **(A)** RFP and **(B)** Optical density measured at 600 nm (OD600) of OptoCre-REDMAP (lacking *ho1-pcyA*) with the plasmid *rfp* reporter containing *ho1-pcyA*. Cells were induced with a range of IPTG concentrations, as indicated, for different light exposure durations. Bars represent the mean values, with individual biological replicates shown as dots (n = 3). A.U., arbitrary units.

**Table S1.** Primers used for colony PCR and qPCR.

| | |
|---|---|
| CCCGATCTTCCCCATCGG | PCR primer to check terminator excision for plasmid *rfp* reporter and chromosomal *rfp* reporter - Forward |
| CTCGAACTCGTGACCGTTAACAG | PCR primer to check for terminator excision for plasmid *rfp* reporter and chromosomal *rfp* reporter - Reverse |
| GAGCACATCAGCAGGACGCACTG | PCR primer to check for gene excision for chromosomal *mcherry-cat* reporter - Forward |
| GGCCCAGTCTTTCGACTGAGCCT | PCR primer to check for gene excision for chromosomal *mcherry-cat* reporter - Reverse |
| AGGGACACGGCGAAATAAC | qPCR primer to check for presence of terminator on plasmid *rfp* reporter - Forward |
| TTTGATGCCTGGAGATCCTTAC | qPCR primer to check for presence of terminator on plasmid *rfp* reporter - Reverse |

**Supplementary Movie Captions**

**Movie S1.** Single-cell dynamics of OptoCre-REDMAP with the plasmid *rfp* reporter in the mother machine microfluidic device with 0 μM IPTG. Red light exposure begins at t = 3 hours.

**Movie S2.** Single-cell dynamics of OptoCre-REDMAP with the plasmid *rfp* reporter in the mother machine microfluidic device with 50 μM IPTG. Red light exposure begins at t = 3 hours.
